# Interaction and co-phase separation of SARS-CoV-2 nucleocapsid protein and human hnRNPA1 and its implications for viral life cycle

**DOI:** 10.1101/2025.11.28.691101

**Authors:** Parul Gupta, Risabh Sahu, Iman Nandi, Subba Rao Gangi Setty, Saumitra Das, Mahavir Singh

## Abstract

The nucleocapsid (N) protein of SARS-CoV-2 is central to viral assembly and replication. It binds viral RNA to form a helical nucleocapsid and enabling genome packaging and release into host cells. Human hnRNPA1, one of the most abundant RNA-binding proteins in eukaryotes, regulates key aspects of RNA metabolism, including splicing, transcription, localisation, and transport. Here, we report a direct physical interaction between SARS-CoV-2 N protein and hnRNPA1 with moderate affinity (Kd ∼ 0.18 μM), primarily mediated through their intrinsically disordered regions. Furthermore, we found that these proteins co-phase separate *in vitro* and co-localise within stress granules in cells. *In vivo* studies reveal that hnRNPA1 suppresses viral replication, suggesting that the N protein – hnRNPA1 interaction plays an important role in modulating the viral life cycle.

## 1 INTRODUCTION

Severe acute respiratory syndrome coronavirus 2 (SARS-CoV-2) is an enveloped, positive-sense single-stranded RNA virus responsible for COVID-19 pandemic^1,2^. Continuous mutations in SARS-CoV-2 raise potential challenges for the efficacy of existing vaccines^3^. Although the vaccines, presently available, are able to contain the impact of the virus, it is important to understand the molecular mechanisms that regulate viral replication, assembly, and interaction with the host cellular machinery for developing targeted therapeutics and vaccines against future outbreaks.

The nucleocapsid (N) protein of SARS-CoV-2 is a multi-functional 419 amino acid (a.a) long structural protein critical for packaging of viral genome and virion assembly ^4–6^ (Figure 1A). N protein is the most abundant protein in coronavirus, and binds viral RNA, forming a ribonucleoprotein (RNP) complex to protect it from the host cell environment ^5–7^. Furthermore, it enhances the efficiency of sub-genomic viral RNA transcription and is essential for viral replication ^6,8,9^. N protein has two conserved folded domains: N-terminal domain (NTD, residues 51-174) and C-terminal domain (CTD, residues 247-365), responsible for RNA binding and dimerisation of protein, respectively (Figure 1A and B). The two folded domains are separated by a Ser/Arg (SR)-rich intrinsically disordered (IDR) linker domain and are flanked on both termini by two additional IDRs, namely N-arm (residues 1-51) and C-tail (residues 365-419)^7,10^ (Figure 1A). Several studies have shown that recombinant N protein can undergo liquid-liquid phase separation (LLPS) *in vitro*^6,11–15^. Upon overexpression of N protein in human cell lines, it is recruited into the stress-granules (SGs)^14,16^. In addition, N protein induces both humoral and cellular immune responses after infection, making it a candidate for antiviral drug therapies^10,17,18^.

**Figure 1.**
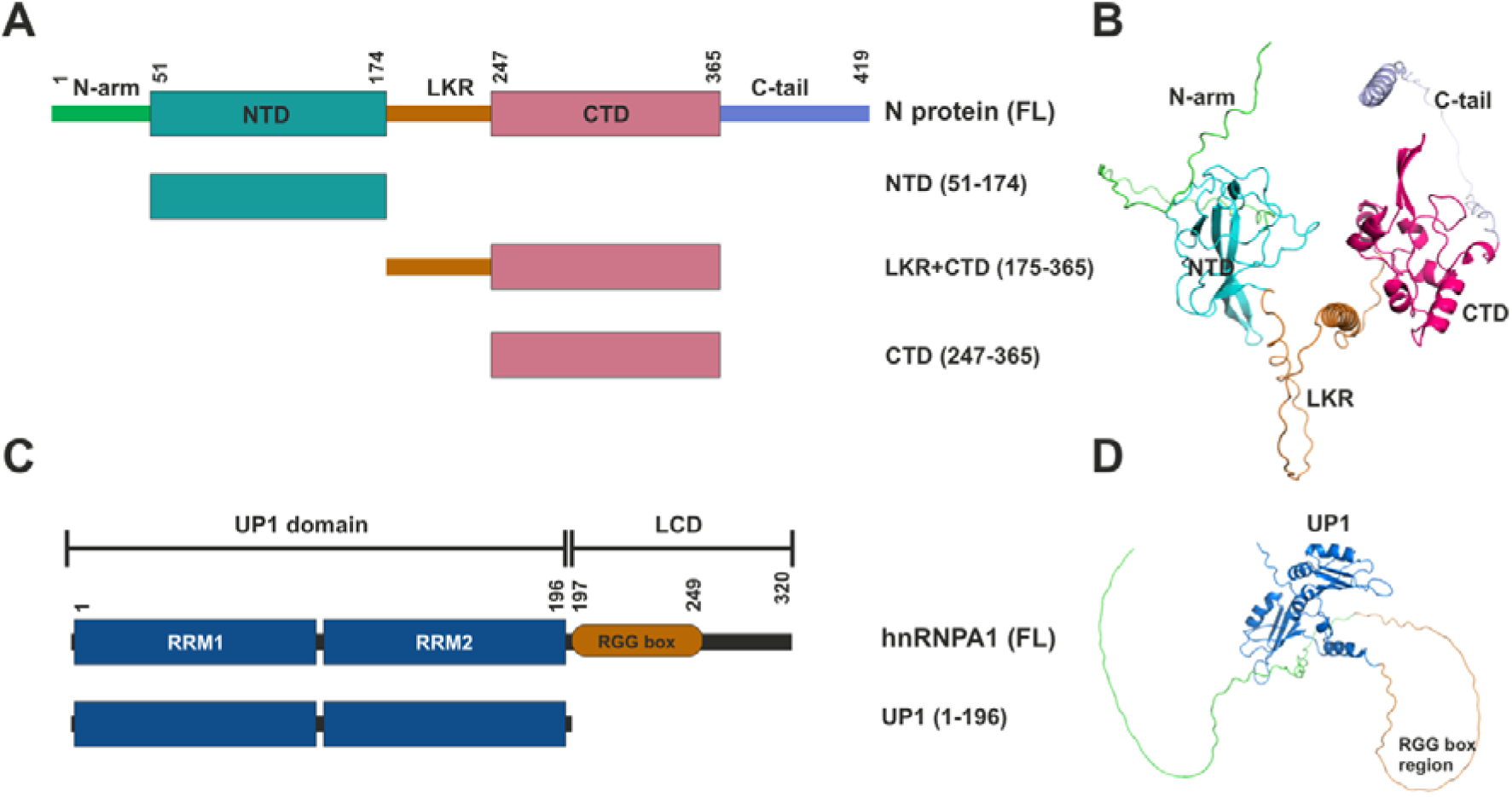
Domain architectures and AlphaFold 2(AF2) predicted structures of full-length SARS-CoV-2 N protein and human hnRNPA1. A) Schematic representation of the full-length N protein and its constructs used in this study. The N protein comprises an N-terminal arm (N-arm), N-terminal domain (NTD, residues 51–174), a central intrinsically disordered linker region (LKR, residues 175–246), C-terminal domain (CTD, residues 247–365), and a C-terminal tail (C-tail, residues 366–419). Truncated constructs used in this study include NTD (51–174), LKR+CTD (175–365), and CTD (247–365), shown below the full-length protein. B) AF2-predicted structural model of the full-length SARS-CoV-2 N protein showing the organisation of individual domains: NTD (cyan), LKR (orange), CTD (magenta), and flexible N- and C-terminal tails (green and light pink, respectively). C) Schematic representation of the human hnRNPA1 protein showing the N-terminal RNA recognition motifs (RRM1 and RRM2) forming the UP1 domain (residues 1–196), followed by the low-complexity domain (LCD; residues 197–320) containing the RGG box (orange). Different constructs used in this study are shown below the full-length protein, along with the domain boundaries. D) AF2-predicted structural model of full-length hnRNPA1 highlighting the UP1 domain (blue) and the disordered low-complexity region containing the RGG box (orange).

hnRNPA1 is 320 residues long, highly ubiquitous protein from the hnRNP A/B subfamily found in the eukaryotic cells^19^. It plays roles in several aspects of RNA metabolism, such as splicing, transport, and localisation of RNA. hnRNPA1 consists of an N-terminal UP1 domain (comprising RRM1 and RRM2 domains, residues 1 to 196) and an intrinsically disordered C-terminal domain, also known as low complexity domain (LCD, residues 197 to 320) (Figure 1C and D). The UP1 domain binds to both single-stranded and double-stranded nucleic acids and its interaction with DNA and RNA have been thoroughly characterized using X-ray crystallography and NMR spectroscopy methods ^20–22^ ^23,24^. UP1 is also a well-established DNA/RNA G-quadruplex destabilising protein^25–29^. The C-terminal LCD of hnRNPA1 contains RGG-box, prion-like domains (PrLD), and nuclear localisation sequence (NLS)^30^. In past decade, there have been several reports that have shown that the RGG-box is a key sensor of G4 structures in the cell^31–33^. The LCD of hnRNPA1 has been shown to drive liquid-liquid phase separation (LLPS) of hnRNPA1^30,33–36^ *in vitro*. The physiological function of hnRNPA1 in the host is defined by its diverse binding affinities to DNA, RNA, and proteins governing processes such as transcription, post-transcriptional modification, translation,, and expression^19,37,38^. Mutations in the hnRNPA1 cause various neurodegenerative diseases such as multisystem proteinopathy (MSP) and amyotrophic lateral sclerosis (ALS)^30,39^.

In a past study, the analysis of SARS-CoV-2 interactome showed that the N protein pulls down hnRNPA1 along with other splicing and stress granule modulators^40^. In another study, using the yeast two-hybrid assay and tandem mass tag-affinity purification mass spectrometry, reported 739 high-confidence protein-protein interactions (PPIs) between 28 viral and 579 human proteins that included the interaction between hnRNPA1 and N protein^41^. hnRNPA1 has been shown to play crucial roles in life cyclic of several viruses. For example, hnRNPA1 has been shown to specifically bind to the 5′ UTR of Sindbis virus (SV) RNA and promotes virus RNA replication and translation^42^. In case of Porcine epidemic diarrhea virus (PEDV), viral nucleocapsid protein and host hnRNPA1 were shown to interact with each other and knockdown of hnRNPA1 expression in CCL-81 cells severely inhibits PEDV replication, suggesting its positive regulatory role in the infection process^43^. In enterovirus 71 (EV71), hnRNPA1 is relocalised to cytosol and promotes internal ribosome entry site (IRES)-mediated translation by interaction of its UP1 domain with conserved stem-loop structures (SLII and SLVI) located within the 5′ UTR^44,45^. Surprisingly, whereas this activity in EV71 is replaceable by hnRNPA2, the homologous cytosolic relocalisation of protein in cells infected with SV that promotes translation in an hnRNPA1-specific fashion^45^. In spite of hnRNPA1’s role in enhancing translation in EV71, siRNA mediated knockdown in cells does not impact EV71 replication to any significant extent, exemplifying its more complex role in its life cycle. Furthermore, hnRNPA1 is an IRES trans-acting factor (ITAF) that binds specifically to 5’ UTR of human rhinovirus-2 and human apoptotic peptidase activating factor 1 (apaf-1) mRNA, thus regulating translation^46^. In addition, in avian reovirus (ARV) infection, hnRNPA1 binds to the p17 protein through its nuclear export signal (NES), driving nucleocytoplasmic shuttling required by the virus to replicate^47^.

In the case of several viral infections, hnRNPA1 functions as a host restriction factor, exerting antiviral effects. For example, in human T-cell lymphotropic virus type I (HTLV-1), hnRNPA1 negatively regulates viral replication by inhibiting the binding of the Rex protein to its response element in the 3′ LTR of viral RNAs^48^. Similarly, during hepatitis C virus (HCV) infection, hnRNPA1 interacts with the 3′ cis-acting replication element (CRE) and the viral RNA-dependent RNA polymerase NS5B, thereby suppressing HCV RNA synthesis. Silencing hnRNPA1 enhances viral replication, suggesting that it competes with the polymerase for binding to the CRE^49,50^. In influenza A virus (IAV), hnRNPA1 knockdown leads to a marked increase in nucleoprotein expression and viral replication, while its overexpression significantly reduces nucleoprotein levels and impairs replication^37^. Collectively, these findings highlight hnRNPA1’s protective, antiviral roles and its potential as a target for host-based antiviral therapeutics^51^.

Interestingly, more than a decade back, there were reports that suggested direct, high-affinity binding between human hnRNPA1 and N protein of Coronaviruses such as mouse hepatitis virus (MHV) and SARS-CoV^52,53^. Here we set out to probe the possible interaction between hnRNPA1 and SARS-CoV-2 N protein to study its implications for viral life cycle. We report that hnRNPA1 and N protein physically interact with moderate affinity. This interaction is mainly driven by the intrinsically disordered regions of the proteins. Furthermore, interestingly, we showed that N protein and hnRNPA1 co-phase separate *in vitro* and colocalizes in the stress-granules in the cells. We have also shown that over-expression hnRNPA1 decreases viral RNA levels and its infectivity in cells, indicating that hnRNPA1 may act as host restriction factor that negatively regulates the SARS-Cov-2 viral life cycle.

## 2 MATERIALS AND METHODS

### Reagents and antibodies

Alexa Fluor 488 C_5_ maleimide (Invitrogen, A10254) and Alexa Fluor 594 C_5_ maleimide (Invitrogen, A10256) were purchased commercially. Commercial antibodies with specific use (IB, immunoblotting; IF, immunofluorescence microscopy; IP, immunoprecipitation; FACS, fluorescence-activated cell sorting) were purchased commercially: Anti-m Nucleocapsid (IB, IF, 40143-MM05), anti-m G3BP1 (IF, sc-365338), anti-r G3BP1 (IF, NBP1-18922), anti-r HA (IB, IF, C29F4). All secondary antibodies were purchased from Invitrogen and used in 1:500 dilutions.

### 2.1 Cloning, protein expression and purification

Plasmids were expressed using *E. coli* strain Rosetta (DE3) and grown on kanamycin and chloramphenicol. Cells are induced with 0.25 mM IPTG after OD_600_ of 0.8 at 37°C. They are further grown at 18°C for 16 hr, centrifuged to obtain cell pellets. Cell pellets were resuspended in buffer A (25 mM Tris–HCl pH 7.5, 5LmM MgCl_2_ 10% glycerol, 5 mM β-mercaptoethanol, 1LmM NaN_3_) plus 500 mM NaCl and 5 mM imidazole pH 8. Cells are lysed by sonication (33% amplitude, 3s on, 6s off) and centrifuged. The supernatant was loaded onto the Ni^2+^-NTA affinity column (Ni-NTA Superflow; Qiagen), washed in buffer A plus 300 mM NaCl and 20 mM imidazole, pH 8.0, and eluted in buffer A plus 300 mM NaCl and 400 mM imidazole. Eluted proteins were concentrated in centrifugal concentrators (Amicon Ultra, EMD Millipore) and further purified by size-exclusion chromatography (Superdex 200/75; GE Life Sciences) under SEC buffer (25 mM Tris–HCl pH 7.5, 300 mM NaCl, 5LmM MgCl_2_, 10% glycerol, 1 mM DTT). SDS-PAGE was used to confirm the presence and purity of proteins. Purified proteins were concentrated and stored at 4°C for further analysis.

### 2.2 Microscale thermophoresis

MST experiments were performed using a Monolith NT.115 (NanoTemper, Munich, Germany) instrument. Protein labelling was carried out with RED-NHS 2^nd^ generation dye from RED-NHS MonolithTM Protein Labeling Kit (NanoTemper Technologies, cat. no.- MO-L011) according to manufacturer’s instructions. 20 µM of protein was incubated with the dye in labeling buffer containing 130 mM NaHCO₃ and 50 mM NaCl at pH 8.2 for 30–60 min at room temperature. Excess unreacted dye was removed using Sephadex G-25 HiTrap desalting columns (Cytiva, Marlborough, MA, USA). Binding titrations were performed by incubating the labelled protein with varying concentrations of ligands. For N protein binding studies, UP1 was used as the ligand. For hnRNPA1 binding assays, NTD, CTD, and LKR+CTD were used as ligands. Ligands were titrated against the labelled protein in a 1:1 dilution series, starting with the highest ligand concentration of 100 μM. A total of 16 two-fold serial dilutions of the target ligand were prepared. The labelled protein was added so that the final concentration of the protein was 5 nM. All the experiments were performed in triplicates for each ligand-protein pair. MonolithTM NT.115 MST Standard Capillaries (NanoTemper Technologies, cat. no.- MO-K022) were used to detect the binding. Curves were fitted with a single-site binding model using PALMIST 1.5.6 software^54^ and figures were generated using GUSSI software v1.1.0^55^.

### 2.3 Nickel-NTA pull-down

Purified 6XHis-N protein (bait) protein was mixed with equal molar ratio of GST-hnRNPA1 (prey) in 500 μl total reaction volume prepared in PBS and 20 mM imidazole pH 8.0. Ni-NTA-agarose beads (30 μl, 50% slurry, Qiagen) were added to protein mixture and incubated for 1 hr at 4°C with occasional mixing. After incubation, beads were washed three times with 2 ml buffer and collected in falcon. The bound proteins were eluted using PBS plus 300 mM imidazole pH 8.0. 10 μl of three collected washes, and each eluate was run on SDS-PAGE for analysis.

### 2.4 Alexa Fluor labelling of proteins

Nucleocapsid protein was labelled with Alexa Fluor 488 C_5_ maleimide (green) and hnRNPA1 was labelled with Alexa Fluor 594 C_5_ maleimide (red). Proteins were labelled with maleimide AlexaFluor dyes by diluting protein stocks into labelling buffer (20 mM HEPES pH 8.3, 300 mM NaCl). Alexa Fluor dyes dissolved in DMSO were added to the protein solutions with a molar ratio of 1:10 of protein and fluorescence dye. 10xTCEP was added to reduce disulfide bonds and then flushed with inert gas. Reactions were incubated for an hour at room temperature and then overnight at 4 ℃ in the dark. Unreacted Alexa Fluor dye was removed next day by buffer exchange using HiTrap desalting columns (Cytiva GE17-1408-01) equilibrated in the labelling buffer. Labelled proteins were then concentrated and buffer exchanged into appropriate storage buffers and flash-frozen.

### 2.5 Phase separation

Labelled N protein and hnRNPA1 are diluted to 50 µM in phase separation buffer (20 mM HEPES, pH 7.4) at varying salt concentrations. For *in vitro* colocalization studies, labelled N protein and hnRNPA1 are mixed in a 1:1 molar ratio in buffer (20 mM HEPES, pH 7.4, 150 mM NaCl). The mixture was incubated for 5 minutes at room temperature. Around 5 µl of solution is transferred to the slide, and images were collected using a Leica SP8 FALCON confocal microscope. ImageJ Fiji was used for image analysis.

### 2.6 Cell culture

HEK-293-T cells expressing ACE-2 (obtained from BEI resources), Vero E6 cells and HeLa (ATCC) cells were maintained in DMEM media supplemented with 10% FBS (Biowest), 1% L-glutamine, and 1% penicillin-streptomycin (Pen-Strep) antibiotics at 5% CO2 in a humified cell culture chamber. Transient transfection of the cells with plasmids was performed post-incubation in OPTI-MEM (Invitrogen) for 30 min and followed the manufacturer’s protocol of Lipofectamine 2000 (Invitrogen). HEK-293-T-ACE-2 cells were used for infection assays and HeLa cells were used for imaging studies.

### 2.7 Immunofluorescence microscopy and image analysis

For steady-state localization studies, cells on coverslips were fixed with 3% formaldehyde (in PBS) and stained with primary antibodies followed by the respective secondary antibodies. Immunofluorescence (IF) microscopy of cells was performed on an Olympus IX81 motorized inverted fluorescence microscope equipped with a CoolSNAP HQ2 (Photometrics) CCD camera using a 60× (oil) U Plan super apochromatic objective. Briefly, cells were imaged for 10–12 z-stacks with a z-step size of 0.2 μm.

The colocalization between two colors was measured by selecting the entire cell and then estimating Pearson’s correlation coefficient (*r*) value using Image J software. The average *r* value from 10 to 20 cells was calculated and then represented as mean ± SEM. Note that the maximum intensity projection of undeconvolved Z-stack images was used to measure *r* values. Final images (maximum intensity projection, MIP) were converted into TIFF format and then assembled using Adobe Photoshop or CorelDraw.

### 2.8 Virus preparation

Virus stocks were procured from BEI resources, NIH, and NIAID and maintained by the viral BSL3 repository at IISc. WT: Isolate Hong Kong/ VM20001061/2020. Vero E6 cells were used for virus propagation and titration, and HEK-293T-ACE-2 cells were used for experiments. MOI of 0.1 was used for virus infection, and cells were harvested after 48 hours.

### 2.9 RNA isolation followed by Real-time PCR

RNA was isolated from HEK-293T-ACE-2 cells using TRI reagent (Sigma) as per manufacturer’s protocol. cDNA was synthesized from RNA using reverse primer for positive strand quantification. M-MLV Reverse transcriptase (Moloney murine leukemia virus) was used to prepare cDNA at 42 °C. Quantitative real-time (qRT-PCR) was performed after cDNA preparation using a DyNAmo HS SYBER green qPCR kit.

### 2.10 Plaque forming assay

Plaque assay was performed in 24 well plates using Vero E6 cells. 1.5×10^5^ cells were seeded in each well of 24 well plates for 90% confluency of cells. The virus-infected supernatant of the previous experiment was collected and serially diluted in multiple dilutions, and those different dilutions of virus-infected supernatant were added in each well of 24 well plates. After 1h of infection at 37 °C, the media was changed with DMEM containing 0.8% agarose and kept for 48h at 37 °C, with a 5% CO_2_ incubator. After 48h, cells were fixed with 4% paraformaldehyde (PFA) and stained with crystal violet for 20 mins. The overlay was removed, and number of plaques was calculated to calculate the virus titre.

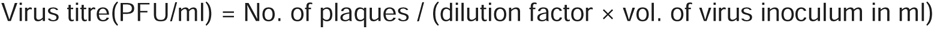

### 2.11 Western blot analysis

The protein concentration of the resulting extracts was determined using the Bradford assay (Bio-Rad). Equal amounts of protein were mixed with SDS sample buffer and heated at 100 °C. Samples were then resolved on 12% SDS–PAGE gels and transferred onto nitrocellulose membranes (NCM, Sigma). Membranes were blocked with 5% skimmed milk at 4 °C on a rocker, followed by overnight incubation with primary antibodies at 4 °C. The next day, membranes were incubated with secondary antibodies for 2 h, and the blots were visualized using ECL substrate (Bio-Rad) on a Chemi-Doc imaging system (Bio-Rad).

## 3 RESULTS AND DISCUSSION

### 3.1 SARS-CoV-2 N protein interacts with hnRNPA1

To probe the direct interaction between hnRNPA1 and SARS-CoV-2 N protein, we first performed a nickel-affinity based pull-down assay. N-terminally GST-tagged hnRNPA1 and N-terminally 6xHis-tagged N protein were overexpressed in *E. coli* and purified separately. Purified N protein was immobilized on the nickel-NTA agarose beads as a bait protein. The GST tagged hnRNPA1 protein acted as the prey protein. After incubation at 4°C and multiple washes, the bound protein was eluted using imidazole in the buffer, and samples were analysed on a SDS-PAGE. As shown in Figure 2A, a distinct band corresponding to hnRNPA1 was detected in the eluted fraction only in the presence of N protein, which was absent in control lanes suggesting that hnRNPA1 physically interacts with N protein.

**Figure 2.**
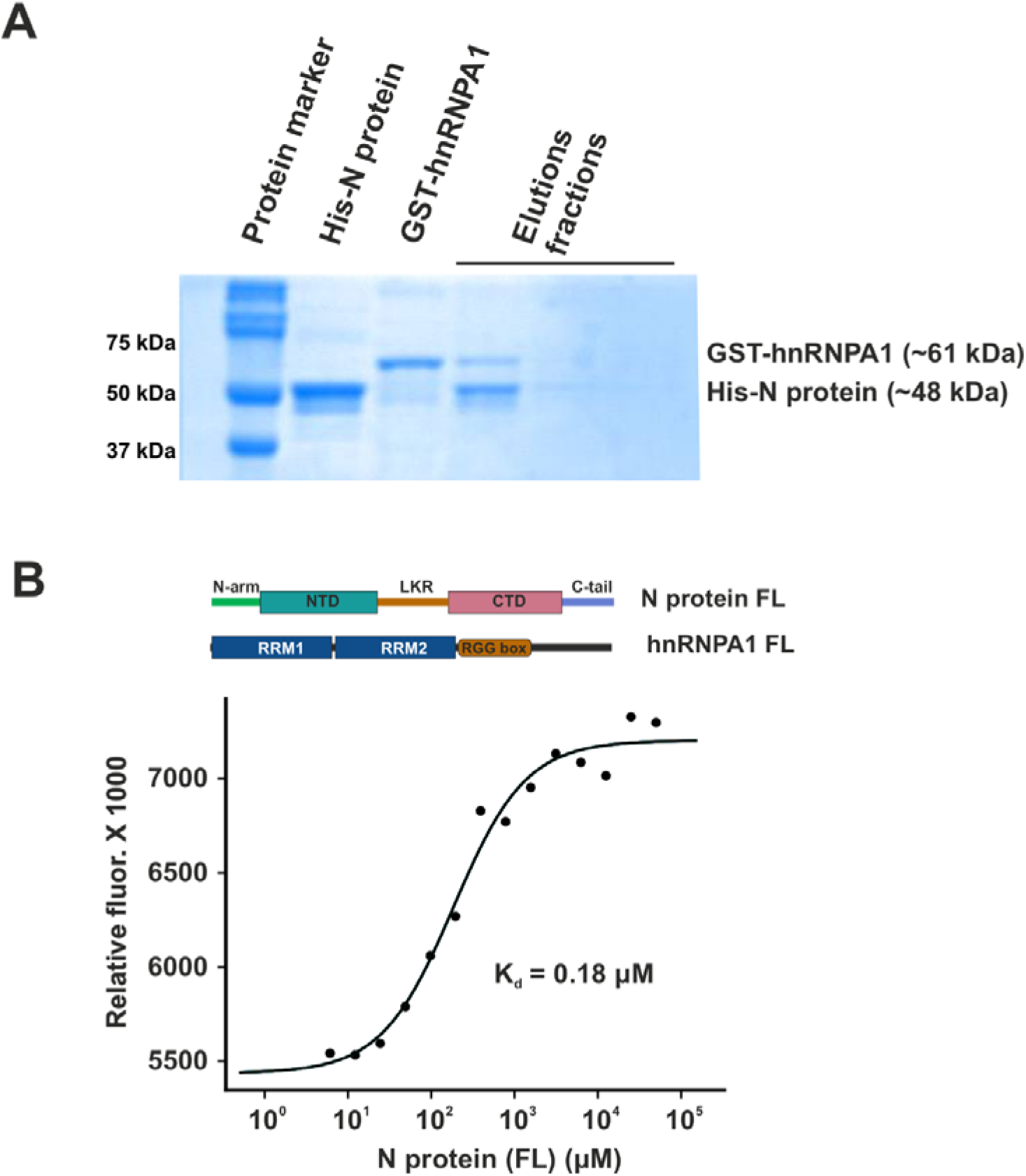
SARS-CoV-2 N protein interacts with hnRNPA1. A) The interaction between SARS-CoV-2 nucleocapsid (N) protein and human hnRNPA1 was determined by Ni²⁺–NTA pull-down assay. Samples were analysed on a 10% SDS-polyacrylamide gel, and protein bands were visualised by Coomassie Brilliant Blue staining. The components loaded in each lane are shown above the gel: lane 1, molecular weight marker; lane 2, purified His-tagged SARS-CoV-2 N protein; lane 3, purified GST-tagged human hnRNPA1; lanes 4–6, eluates showing N protein and pull-down. B) The binding affinity between N protein and hnRNPA1 was determined using MST. The dissociation constant (K_d_) was determined to be approximately 0.18 µM, based on three independent experiments (n = 3).

This interaction was also probed using microscale thermophoresis (MST). MST is a sensitive biophysical technique that measures changes in the mobility of fluorescently labelled molecules under a temperature gradient. Fluorescently labelled hnRNPA1 was titrated with increasing concentration of purified N protein. The MST binding isotherms showed a dose-dependent response consistent with a specific interaction, which was fitted to a one-site binding model. The N protein interaction with hnRNPA1 resulted with a dissociation constant (K_d_) of ∼0.18 µM (Figure 2B).

### 3.2 N protein and hnRNPA1 bind mainly using the disordered domains

Both N protein and hnRNPA1 are modular proteins comprising both the folded domains and disordered regions (Figure 1A-D). To decipher the contribution of the individual domains or the regions for interaction, we purified different domains/regions of N protein and hnRNPA1. This included three constructs of N protein: (i) full-length (FL) N protein (residues 1 to 419), (i) the RNA-binding domain (NTD, also referred to as RBD, residues 51 to 174), (ii) the linker plus dimerization domain (LKR+CTD, corresponding to the central serine/arginine–rich disordered region plus the C-terminal dimerization fold, residues 175 to 365), and (iii) the isolated dimerization domain (CTD, residues 247 to 365) (Figure 1A and Supplementary Figure S1A-D). We also purified full-length hnRNPA1 (residues 1 to 320) and UP1 domain (residues 1 to 196) of hnRNPA1 (Figure 1C and Supplementary Figure S1E-H). As detailed in the previous section, full length hnRNPA1 interacts with N protein with a K_d_ of ∼0.18 µM as determined using MST experiments (Figure 2B). On the other hand, folded UP1 domain of hnRNPA1 did not show a measurable binding with the full length N protein, suggesting that the C-terminal LCD of hnRNA1 mainly binds to the N protein (Figure 3A).

**Figure 3.**
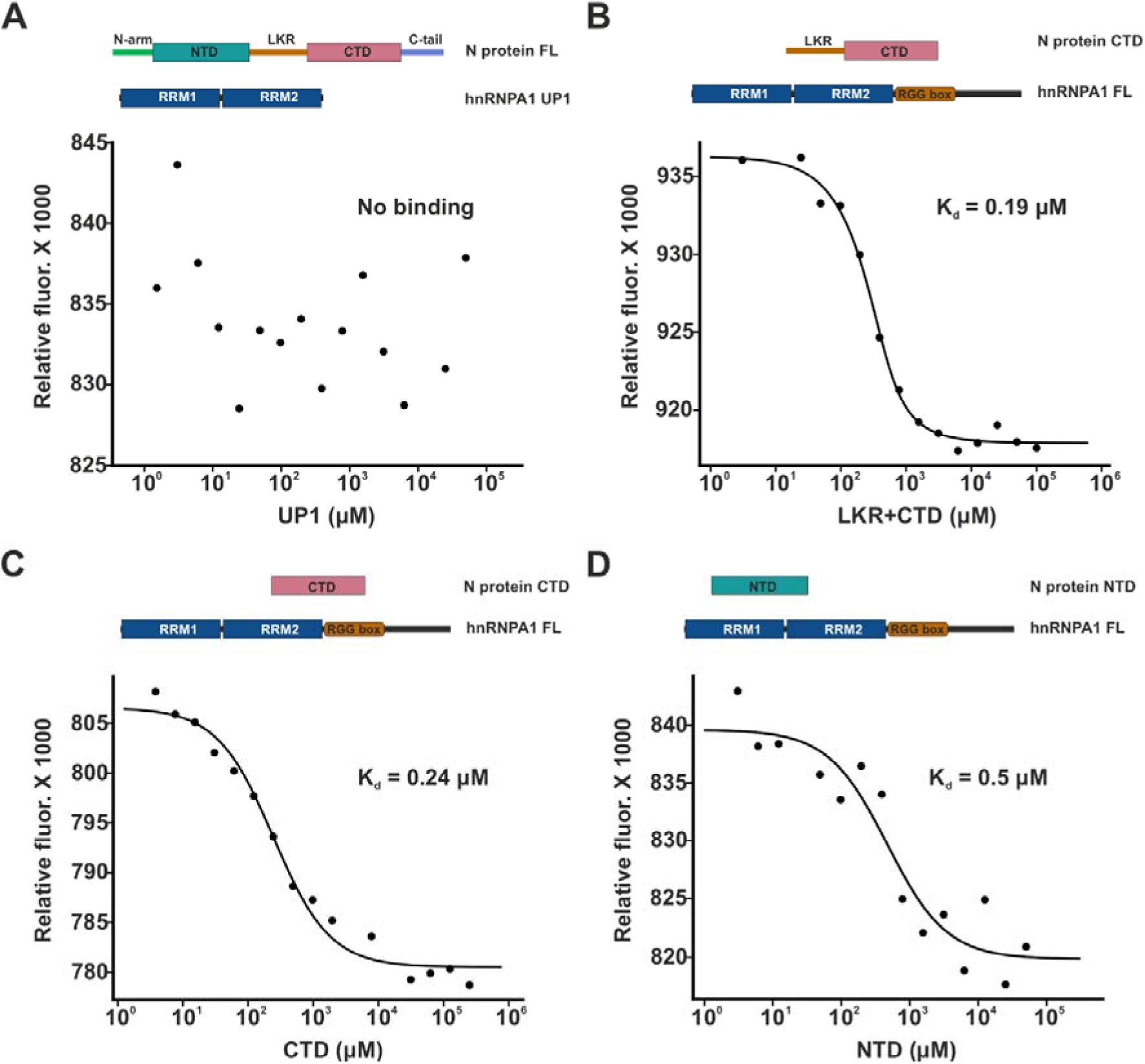
Mapping the interacting domains of SARS-CoV-2 N protein and hnRNPA1. Constructs used in the experiment are shown at the top of each panel. A) MST binding curve showing no detectable interaction between full-length N protein and the UP1 domain (residues 1–196) of hnRNPA1. (B–D) Representative MST binding isotherms depicting the interaction of individual domains of the N protein with hnRNPA1: LKR+CTD (K_d_ = 0.19 µM) (B); CTD (K_d_ = 0.24 µM) (C); NTD (K_d_ = 0.5 µM) (D).

Next, we tested the binding of different domains of N protein for full-length hnRNPA1. All three tested constructs showed measurable binding to hnRNPA1 but with varying affinities. The affinity of hnRNPA1 for LKR+CTD was highest (Kd ∼ 0.19 µM) (Figure 3B) followed by CTD alone (Kd ∼0.24 µM) (Figure 3C), which was followed by NTD alone (0.5 µM) (Figure 3D). Taken together, these results suggest that the NTD-linker-CTD region of N protein binds to the C-terminal LCD region of hnRNPA1.

### 3.3 N protein and hnRNPA1 co-phase separate *in vitro*

Phase separation of viral and host proteins is known to play key role in viral replication and host antiviral defence. N protein has been shown to co-phase separate with proteins of the hnRNP family like FUS and hnRNPA2^56^. Therefore, we tested whether hnRNPA1 and N protein could co-phase separate *in vitro*. First, we determined the droplet formation of both the proteins under different salt conditions (Figure 4A and B). Both hnRNPA1 (Figure 4B) and N protein (Figure 4A) formed spherical, dynamic droplets in buffer with different concentration of salt (NaCl) as observed by confocal microscopy. The number and size of droplets formed by either protein decreased progressively with increasing concentrations of NaCl. At higher salt concentrations (above 100 mM NaCl), the droplets of both hnRNPA1 and N protein were dissolved which confirms that their self-association is sensitive to the ionic strength of buffer and is mediated by charge-charge interactions.

**Figure 4.**
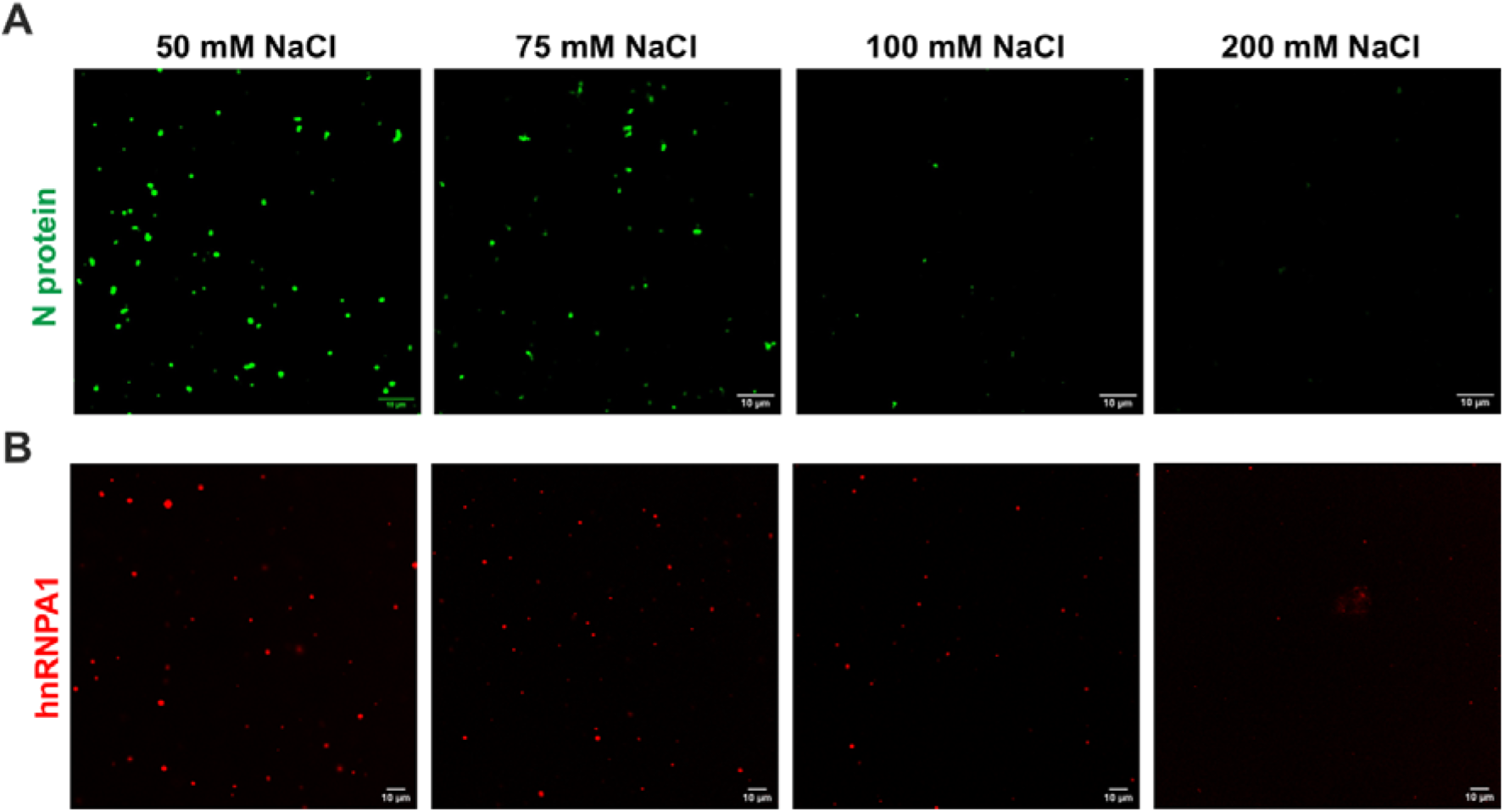
In vitro liquid-liquid phase separation (LLPS) of N protein and hnRNPA1 and its dependence of increasing salt (NaCl) concentration. A) Representative confocal microscopy images showing LLPS of 50 µM A) N protein under varying NaCl concentrations. B) Representative confocal microscopy images showing LLPS of 50 µM A) hnRNPA1 under varying sodium chloride concentrations. The number of droplets decreases with increasing salt concentration from 50 mM NaCl to 200 mM NaCl in both the cases. Scale bars, 10 µm.

Based on the above observation we chose a condition (150 mM NaCl) where N protein and hnRNPA1 formed very few number of liquid-liquid droplets. We used fluorophore-labelled hnRNPA1 (hnRNPA1-Alexa 594) and N protein (N-Alexa 488) proteins in these experiments. In the control experiments, when either hnRNPA1 or N protein was mixed with BSA, we did not observe droplet formation, as observed under a brightfield microscope (Figure 5A, left and middle panels). Interestingly, however, under the same condition, when hnRNPA1 and N protein were mixed together and visualised under brightfield microscope, we observed significant increase in the formation of droplets (Figure 5A, right panel), also reaffirming the specificity of N protein and hnRNPA1 interaction. The co-assembled droplets displayed characteristic liquid-like properties, including spherical morphology, fusion events, and dynamic rearrangement, suggesting that the heterotypic interaction between hnRNPA1 and N protein enhances phase separation. This observation indicates that direct protein–protein interaction can overcome the inhibitory effect of ionic strength to promote droplet formation.

**Figure 5.**
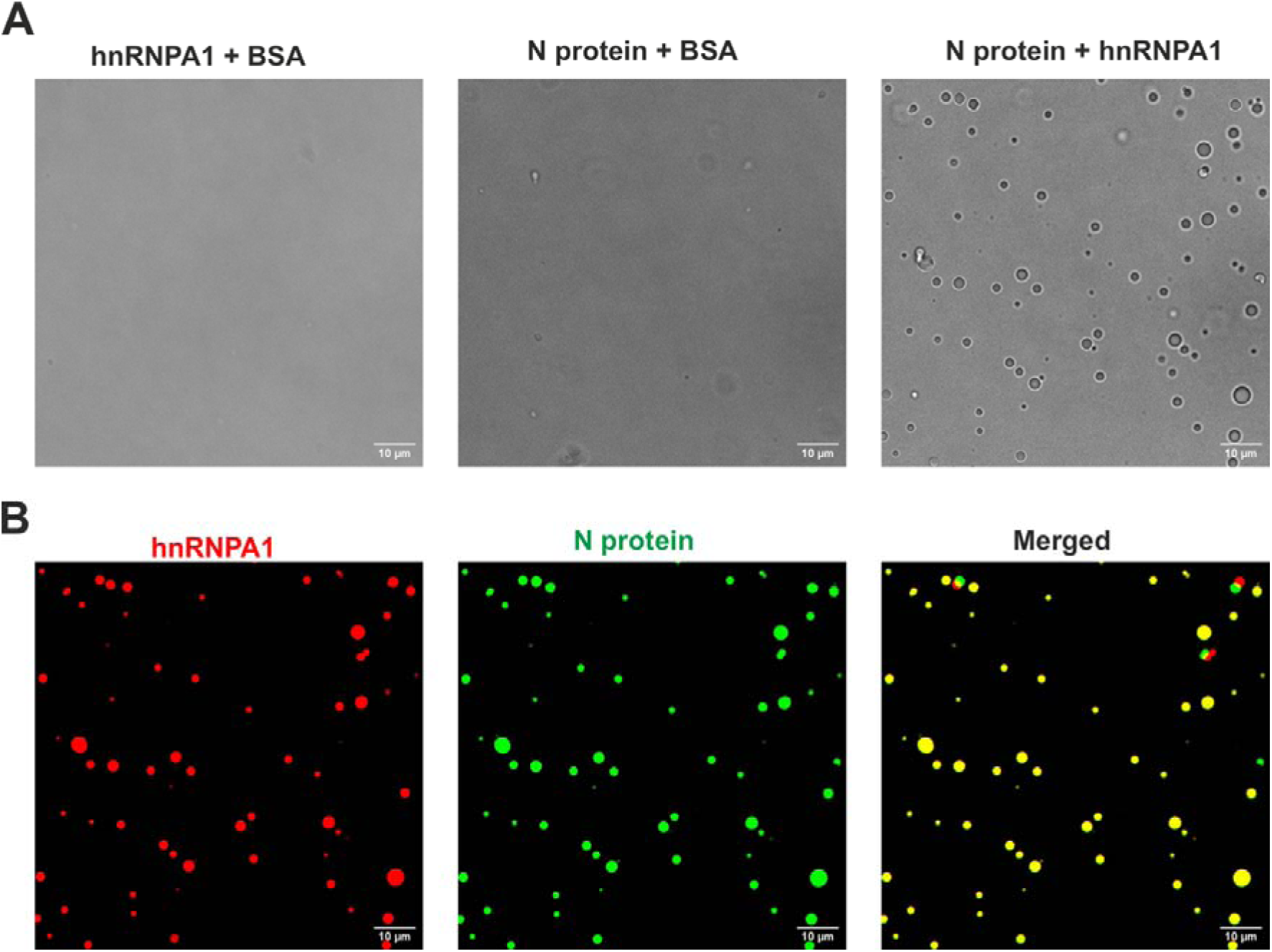
HnRNPA1 and N protein co-phase separate in vitro. A) Differential interference contrast (DIC) images showing that neither hnRNPA1 nor N protein formed droplets when incubated individually with BSA (control). In contrast, mixing hnRNPA1 with N protein led to the formation of droplets. B) Confocal fluorescence images showing co-phase separation of fluorescently labelled N protein (green) and hnRNPA1 (red) within the spherical condensates. The merged image panel displays yellow droplets, indicating that both proteins coexist within the same phase-separated assemblies. Images were acquired at a protein concentration of 50 µM in a buffer containing 25 mM HEPES (pH 7.4), 150 mM NaCl, and 2 mM DTT. Scale bars, 10 µm.

Next, we visualized the co-assembled droplets formed by hnRNPA1-Alexa Fluor 594 and N-Alexa Fluor 488 at 150 mM NaCl conditions using confocal microscopy (Figure 5B). In most of the droplets, both N protein and hnRNPA1 were visualised. These results confirm that hnRNPA1 co-phase separates with N protein *in vitro*, rather than the liquid-liquid phase separation of one protein being promoted by the other. The ability of hnRNPA1 to nucleate N protein into droplets may also mimic the cellular context such as stress granules or other ribonucleoprotein condensates to provide a scaffold for host proteins to trap viral components. Overall, these findings reveal that hnRNPA1 not only binds N protein directly but also promotes its co-assembly into dynamic condensates.

### 3.4 N protein and hnRNPA1 colocalise in the cells

Several past studies have reported that both N protein and hnRNPA1 form biomolecular condensates inside cells and are recruited into the stress granules upon stress^34,57,58^. Also, the propensity of N protein for liquid–liquid phase separation increases with RNA and using this property it can partition into condensates formed by host RNA-binding proteins (hnRNP family members, TDP-43, FUS, etc.) inside the host cell^59^.

To assess whether N protein and hnRNPA1 colocalise within stress granules, HeLa cells were cotransfected with mammalian expression plasmids encoding N protein or hnRNPA1. After 24 h, oxidative stress was induced using sodium arsenite (0.1 mM, 60 min), a known inducer of stress granule formation. Cells were fixed, immunostained with antibodies against N protein, hnRNPA1, and the stress granule marker G3BP1, and visualized by fluorescence microscopy. Their localisation was determined under control and stress environment in HeLa cells. Under control (PBS-treated) conditions, representative confocal images showed that both N protein and hnRNPA1 were diffused within the cell. N protein was spread throughout the cytosol, whereas hnRNPA1 was mainly observed in the nucleus with a faint cytosolic signal, and neither protein formed any visible puncta (Figure 6A). Accordingly, the merged images showed very little spatial overlap between the two proteins. Upon stress (sodium arsenite treatment), both proteins show redistribution within the cell. N protein assembled into distinct cytoplasmic puncta and hnRNPA1 translocates from nucleus to cytoplasm to form bright granular structures (Figure 6B). In the merged images, these puncta showed co-recruitment and clear colocalization, which became even more apparent in the insets highlighting their presence within the same cytoplasmic foci. To verify that these structures were stress granules (SGs), we co-stained the cells with the stress-granule marker G3BP1. Under non-stressed conditions, neither N protein nor hnRNPA1 formed detectable granules and remained diffused throughout the cell. However, upon stress induction, both proteins localized to G3BP1-positive stress granules (Supplementary Figure S2A and B). Quantitative analysis revealed a high degree of signal overlap, indicating direct recruitment of the viral N protein into host ribonucleoprotein condensates (Supplementary Figure S2C).

**Figure 6.**
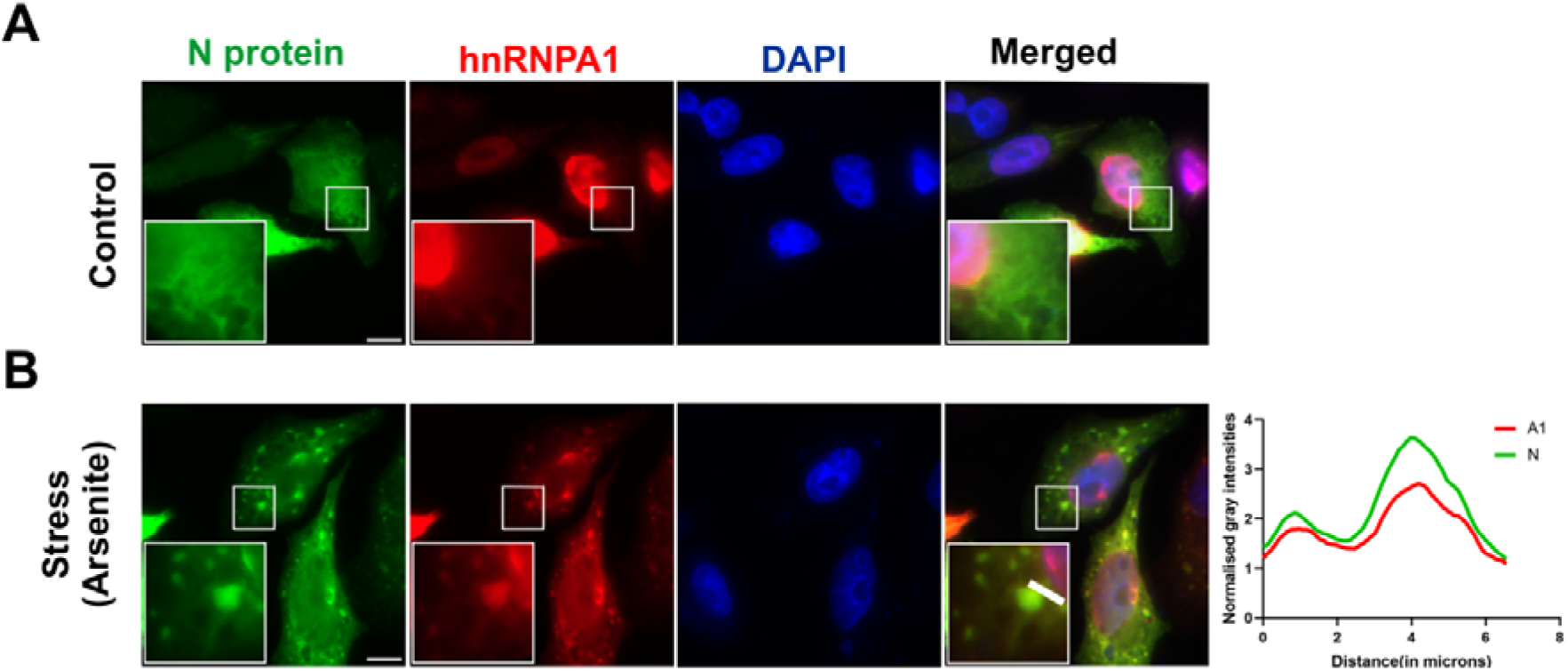
Immunofluorescence images showing colocalization of N protein with hnRNPA1 upon inducing stress. A) Representative confocal fluorescence images of HeLa cells transfected with SARS-CoV-2 N and HA-hnRNPA1 under control (PBS-treated) conditions. B) Confocal images of cells treated with 500 µM sodium arsenite for 1 hr, showing enhanced punctate colocalization of nucleocapsid and hnRNPA1. After fixation, cells were immunostained for N protein (green), HA (red), and nuclei (DAPI; blue). Merged images show colocalization of N protein and hnRNPA1 within the cells. Magnified insets show enlarged views of boxed regions, highlighting representative sites of protein colocalization. Scale bars, 10 µm. Line-scan fluorescence intensity profiles showing normalised signal distributions of hnRNPA1 (A1) and Nucleocapsid (N) proteins along selected transects through representative granules. Overlapping peaks between A1 and N protein indicate their spatial co-enrichment within stress granules.

### 3.5 Overexpression of hnRNPA1 affects the virus infectivity

Based on our findings that N protein and hnRNPA1 interact and colocalize in the same cellular compartment during stress, we wanted to know if SARS-CoV-2 infection is influenced by hnRNPA1 protein expression. For this, hnRNPA1 was overexpressed (cell transfected with human hnRNPA1 cloned in a mammalian expression vector, pcDNA3.1-hnRNPA1), in HEK293-ACE2 cells, followed by infection with SARS-CoV-2 virus (MOI = 0.1) and then subjected to western blot. Briefly, HEK293-ACE2 cells were transfected with either pcDNA3.1 control plasmid or pcDNA3.1-hnRNPA1 plasmid, followed by SARS-CoV-2 infection. Cells were harvested at 24 h post-infection for further analysis (Figure 7A). Virus mRNA levels were studied via RT-PCR under the influence of hnRNPA1 overexpression in SARS-CoV-2 infected cells. Cells with overexpressed hnRNPA1 showed a significant decline (∼42%) in viral mRNA levels (n = 3), p < 0.05 in comparison to pcDNA3.1 transfected cells (negative control), suggesting an inhibitory role of hnRNPA1 at the transcription level in SARS-CoV-2 infection (Figure 7B). Furthermore, a plaque assay was performed to determine the virus infectivity upon hnRNPA1 overexpression. In agreement with the RT-PCR data, a significant reduction in plaque numbers (∼2.5 fold) was observed in hnRNPA1 overexpressed cells compared to the control indicating that not only viral RNA synthesis but also the generation of infectious progeny virus is impaired (Figure 7C and D). This suggests that hnRNPA1 impairs viral life cycle and plays a protective role in SARS-CoV-2 infection. Given the importance of N protein for efficient viral RNA synthesis, protection, packaging, even the partial entrapment of N protein by hnRNPA1 would result in observable reduction in viral RNA and infectivity, as observed in our experiments.

**Figure 7.**
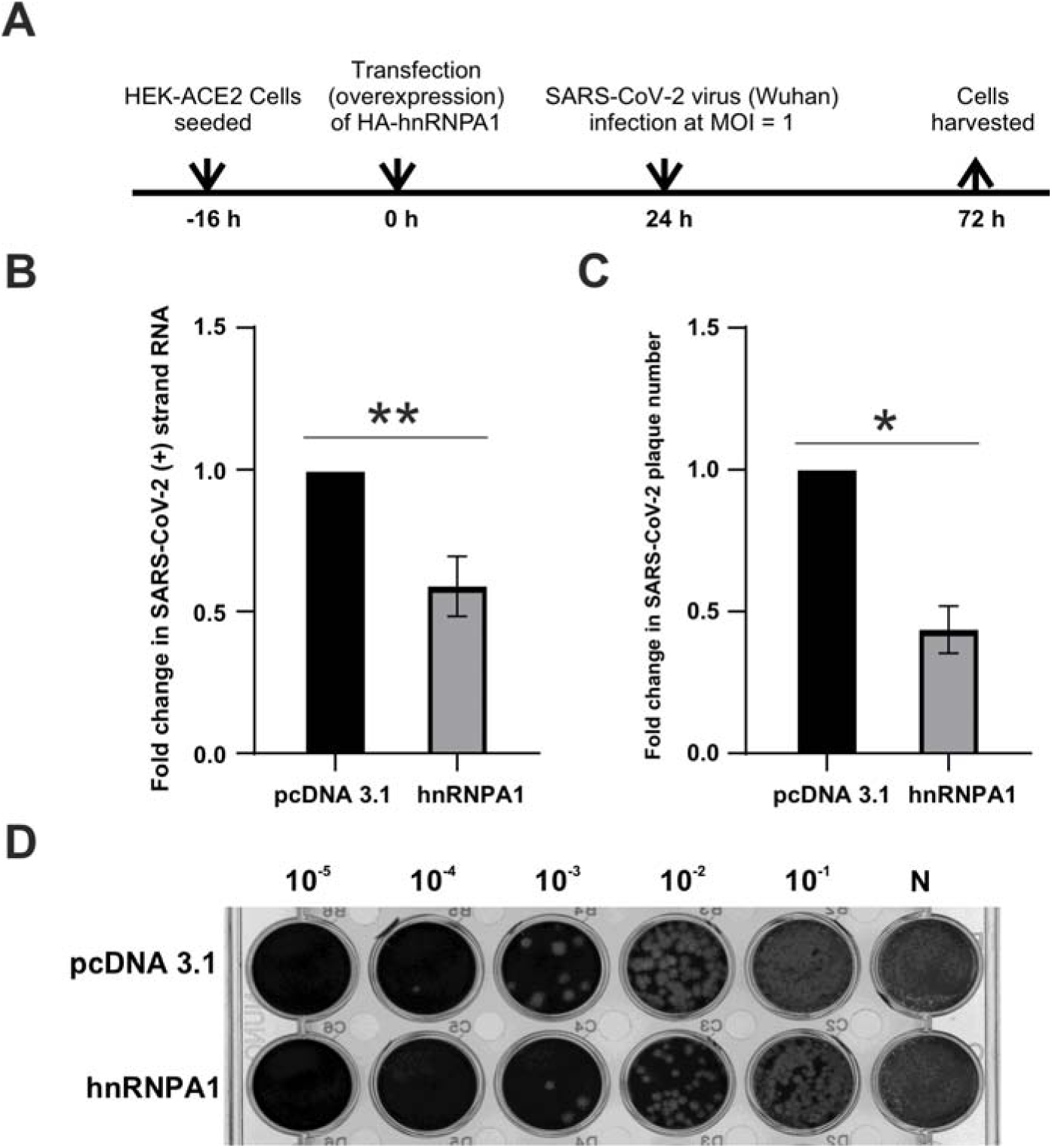
Overexpression of hnRNPA1 reduces SARS-CoV-2 replication in HEK-ACE2 cells. A) Schematic diagram illustrating the experimental workflow. HEK-ACE2 cells were seeded 16 h prior to transfection with either empty vector (pcDNA 3.1) or HA-tagged hnRNPA1 expression construct. After 24 h of transfection, cells were infected with SARS-CoV-2 (Wuhan strain) at a multiplicity of infection (MOI) of 1. Cells were harvested 48 h post-infection (72 h post-seeding) for viral RNA quantification and plaque assays. Arrows indicate key experimental steps during the 72 hr timeline. B) Quantitative RT-PCR analysis showing the relative fold change in positive-sense SARS-CoV-2 genomic RNA compared with the pcDNA3.1 control. hnRNPA1 overexpression led to a significant reduction in viral RNA abundance. Data are presented as mean ± SEM from n = 3 independent experiments (P < 0.01). C) Quantification of infectious SARS-CoV-2 particles by plaque assay. Viral titres are shown as fold change relative to pcDNA 3.1-transfected (control) cells. hnRNPA1 expression significantly decreased the production of infectious viral particles. Data represent mean ± SEM from n = 3 biological replicates (P< 0.05). D) Representative plaque assay images showing a dose-dependent reduction in plaque numbers in hnRNPA1-overexpressing cells compared to the vector control. Serial dilutions (10⁻□ to 10⁻¹) of the culture supernatants were used to infect Vero E6 cells, followed by crystal violet staining to visualise plaques.

## 4 DISCUSSION

hnRNPA1 is ubiquitously expressed nucleocytoplasmic shutting protein inside eukaryotic cells that has various functions in several RNA metabolism events such as pre-mRNA splicing, RNA trafficking as well as in telomere maintenance^60,61^. hnRNPA1 has two RNA recognition motifs (RRM1 and RRM2, together making UP1 domain) involved in RNA binding and a C-terminal glycine rich low complexity domain that promotes liquid–liquid phase separation (LLPS)^34,60^. Under stress, hnRNPA1 translocates from nucleus to the cytoplasm and gets accumulated inside the stress granules (SGs)^62^. In some cases, viral infection also induces formation of cytoplasmic condensates similar to the stress-granules that are formed under various cellular stresses such as heat, oxidation, hypoxia and osmotic pressure. Stress granules are dynamic condensates that are rich in stalled translation initiation complexes, mRNA, and RNA-binding proteins such as G3BP1, TIA-1/TIAR, etc. SGs help cells sorting mRNAs—either storing them for later translation or directing them for degradation in processing bodies (P-bodies)^63–65^.

Many viruses have evolved strategies to manipulate SGs. Enteroviruses cleave G3BP1 to prevent SG assembly^66^, while alphaviruses sequester G3BP1 into replication complexes, thus preventing SG-mediated antiviral responses^67^. Coronaviruses also modulate SGs. For example, in SARS-CoV-2, the N protein remodels SGs by directly interacting with G3BP1/2^14,6814,68,69^. Early in the COVID-19 pandemic, protein–protein interaction studies revealed that the N protein interacts with the stress granule component G3BP1/2. This interaction depends on an ITFG motif located within the N-terminal intrinsically disordered region (N-arm) of N protein, spanning residues R14–G18 (RITFG). This motif interacts with the NTF2-like domain of G3BP^70^. To disrupt this interaction, Long et al. generated a mutant (named the RATA mutant), in which I15 and F17 were substituted with alanine (I15A/F17A). This double mutation abolishes the N–G3BP interaction, thereby preventing G3BP recruitment to viral replication sites and impairing the suppression of stress granule formation during infection which lead to reduced viral replication both in *in vitro* and in animal models^71^. These observations indicate that intact SGs exert an antiviral response, whereas N protein acts as antagonist of virus and promote virus for replication.

In this context, our findings that hnRNPA1 directly binds to N protein and co-phase separates with N protein *in vitro* and inside cells into SGs and inhibition of viral replication upon overexpression of hnRNPA1 poses hnRNPA1 as novel host restriction factor. These findings support previous reports that showed that N protein partitions into condensates along with RNA-binding proteins and suggest that hnRNPA1 can redirect N protein into non-productive assmeblies^58,59^.

There are several plausible mechanisms which may contribute for the observed antiviral activity of hnRNPA1. Firstly, sequestration of N protein into antiviral condensates is one likely mechanism by which hnRNPA1 could play inhibitory role for SARS-CoV-2 virus. By directly interacting with N protein and recruiting it into the stress granules, hnRNPA1 could reduce the pool of available N protein thus affecting viral replication/transcription or genome packaging into virions. Since, N protein is crucial for viral RNA stability and transcription regulation, its sequestration into the stress granules would compromise viral life cycle^13^. Second scenario is that hnRNPA1 may compete with the N protein for viral RNA binding. Both hnRNPA1 and N protein are well recognized RNA binding proteins^59,60^. In the cell, hnRNPA1 might outcompete N protein for viral RNA or sequester viral RNA into non-reproductive ribonucleoprotein complexes. This would reduce the N protein’s ability to stabilise and regulate the viral genome and regulate transcription resulting in virus reduction. In the third scenario,hnRNPA1 directly promotes stress granule stabilization. SARS-CoV-2 N protein has been reported to inhibit SG assembly through interactions with G3BP1/2, modulating the host antiviral response^14,68^. It is possible that overexpression of hnRNPA1 shifts the equilibrium towards SGs persistence, thus enhancing antiviral response. Previous reports suggest that mature SGs not only sequester viral proteins but also contains proteins responsible for activation of innate immunity inside host, hence, reducing viral protein synthesis. Therefore, hnRNPA1 can indirectly enhance interferon response by promoting SGs prolongation.

It is noteworthy that hnRNPA1’s role in viral infections is highly context dependent. While we demonstrate an antiviral function against SARS-CoV-2, hnRNPA1 has been described as a proviral factor in other RNA virus infections. For instance, hnRNPA1 has been shown to promote hepatitis C virus replication and acts as an internal ribosome entry site (IRES) trans-acting factor for enteroviruses^72,73^. On the other hand, hnRNPA1 overexpression restricts replication of vesiculoviruses^74^. In HIV-1, hnRNPA1 regulates viral RNA splicing in ways that can either support or inhibit productive infection. Monette et al. reported that the levels of hnRNPA1 increased post HIV-1 infection and found co-immunoprecipitation of viral RNA and hnRNPA1^75^ When hnRNPA1 levels were reduced, the production of the viral structural protein pr55Gag decreased, leading to reduced viral replication. Additionally, HIV-1 infection was found to interfere with the import of hnRNPA1 into the nucleus. Cytoplasmic retention of hnRNPA1 favoured IRES-mediated translation of viral genes and vice-versa^51,75–77^. In contrast, Zahler et al highlighted the importance of hnRNPA1 in regulating the splicing of many viral genes (viz, Tat), wherein, hnRNPA1 binds to Exonic Splicing Silencer 2 (ESS 2), thereby repressing the splicing of Tat mRNA. Overexpression of hnRNPA1 in vitro significantly reduced Tat protein levels and consequently impaired viral replication^51,78–80^. This suggests the dual, proviral or antiviral nature of hnRNPA1 and other RNA binding proteins. Their activity is context dependent determined by the virus-host interactions.

Therefore, we suggest a model wherein hnRNPA1 sequesters SARS-CoV-2 N protein into stress granules by direct protein–protein interactions, leading to reduced viral RNA metabolism, nucleocapsid assembly, and replication. The contrasting literature, where N antagonizes SG formation but hnRNPA1 overexpression restores SG persistence-suggests a dynamic tug-of-war. The outcome may depend on the relative stoichiometry of viral N protein and host RBPs. In conclusion, our data identifies hnRNPA1 as multifunctional host restriction factor against SARS-CoV-2. By directly binding N protein and redirecting it into stress granules, hnRNPA1 limits viral RNA metabolism, nucleocapsid assembly, and replication and can be explored as a potential therapeutic target by exploring its modulation.

## Data availability statement

The data underlying this article are available in the article and in its online supplementary material.

## Supplementary data

Supplementary data are provided in the SI file.

## Acknowledgements

The authors acknowledge funding for infrastructural support from the following programs of the Government of India: DST FIST, DST, DBT SAIF, UGC CAS, and the DBT-IISc partnership program. We acknowledge the divisional bioimaging facility for imaging.

## Author contribution

M.S. conceived the research and acquired the funds. P.G. designed and carried out the experimental design and performed cloning, protein purification, microscale thermophoresis, pull-down assays, fluorescent labelling, and phase-separation experiments. R.S. conducted the viral infection studies, RT-PCR, and plaque assays. P.G. and I.N. performed cell culture work and immunofluorescence experiments. S.D. and S.R.G.S. provided reagents and helped in experiment design. Data analysis was performed by P.G. and M.S. The manuscript was written by M.S. and P.G. with the input from all the authors.

## Funding

Authors acknowledge funding support from the Anusandhan National Research Foundation (ANRF), Department of Science and Technology (DST), India (STAR award number STR/2021/000015 and CRG grant number CRG/2021/003597 to M.S.) and Ministry of Education (MoE). This work was also supported by the Department of Biotechnology (BT/PR32489/BRB/10/1786/2019) and Science and Engineering Research Board (CRG/2019/000281) to S.R.G.S.

## Conflict of interest statement

None declared.

## Notes

### Competing Interest Statement

The authors have declared no competing interest.

