## Supplemental Figures for "Interaction and co-phase separation of SARS-CoV-2 nucleocapsid protein and human hnRNPA1 and its implications for viral life cycle"

This file contains:

1. Supplementary Figure S1

2. Supplementary Figure S2

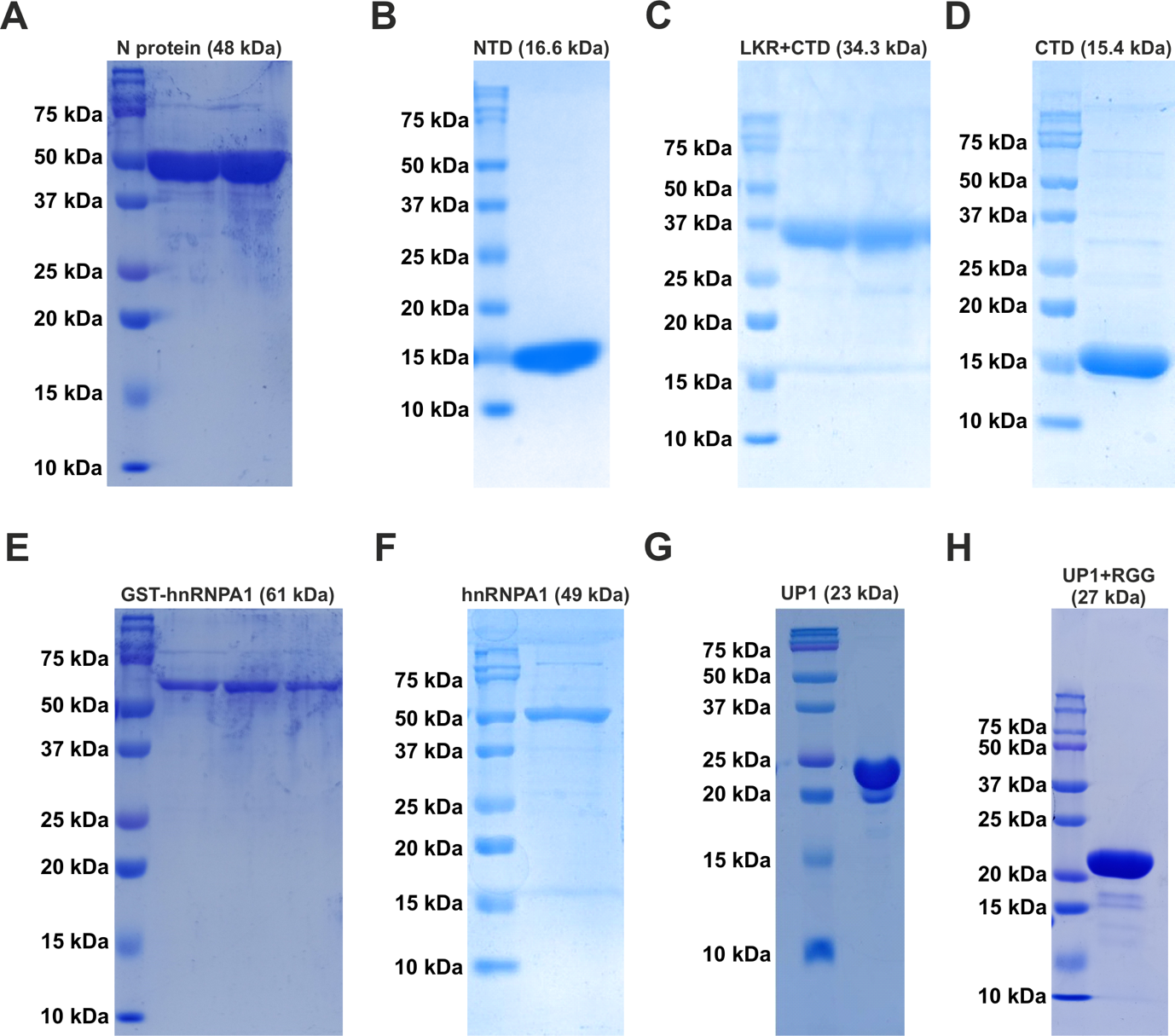

**Supplementary Figure S1.** SDS-PAGE images of purified proteins used in this study. A-D) Purification of hnRNPA1 and its truncated constructs. Left to right: GST-hnRNPA1 fusion protein (61 kDa) (A); Full-length hnRNPA1 (49 kDa) (B); UP1 domain (23 kDa) (C); UP1+RGG fusion (27 kDa) (D). E-H) Purification of N protein and its truncated constructs. Left to right: Full-length N protein (48 kDa) (E); NTD (16.6 kDa) (F); LKR+CTD fragment (34.3 kDa) (G); CTD (15.4 kDa) (H).

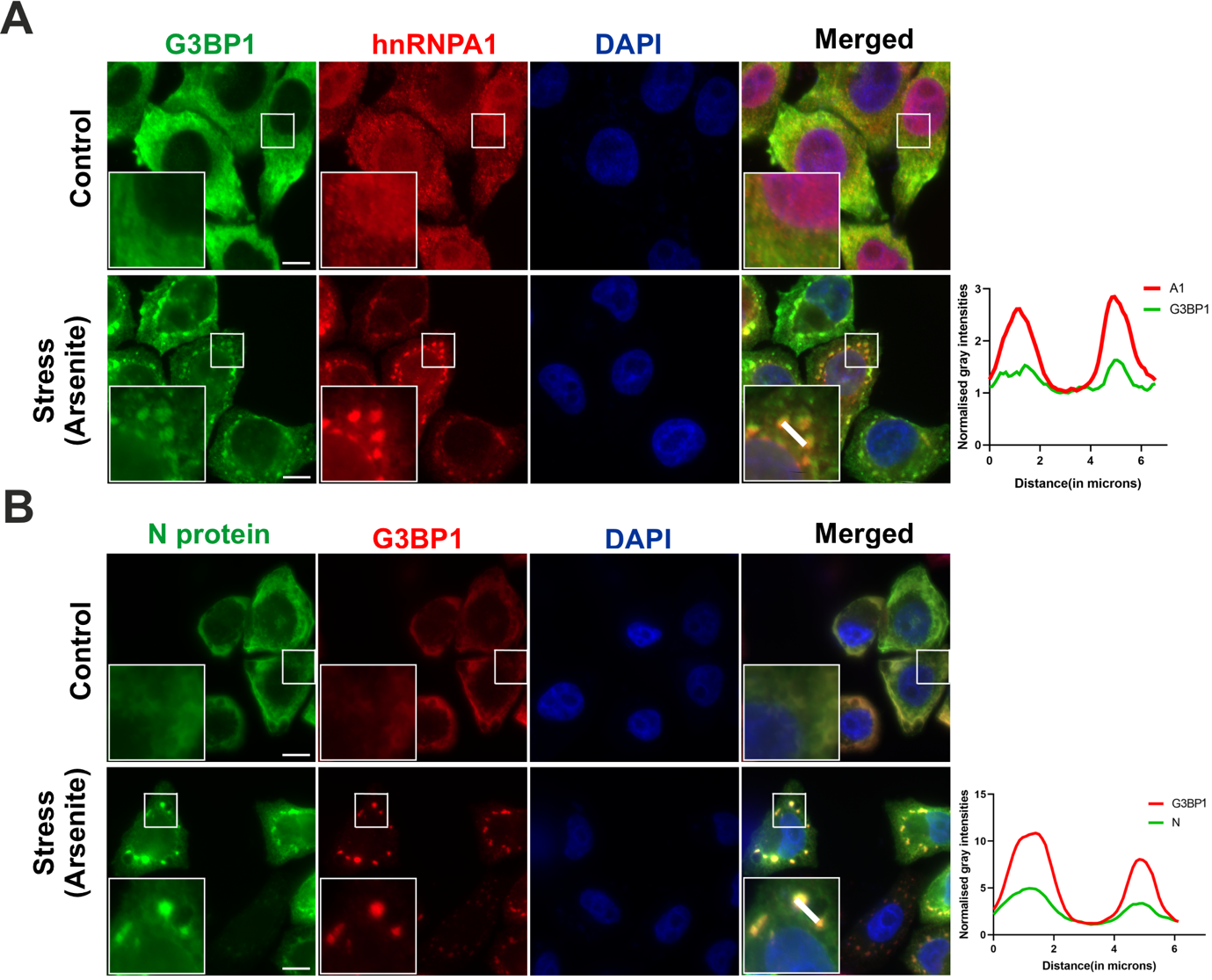

**Supplementary Figure S2.** Localization of N protein and hnRNPA1 upon stress to the stress-granules (SGs). Cells were transfected with the indicated constructs and treated with 500 μM sodium arsenite for 1 hr to induce stress granules. A) Representative confocal images showing colocalisation of hnRNPA1 protein with G3BP1 in HeLa cells. Cells were stained with DAPI, anti-HA and anti-G3BP1. hnRNPA1 is present in nucleus (control) which gets translocated to SGs upon stress. Line-scan fluorescence intensity profiles showing normalised signal distributions of G3BP1 and NhnRNPA1 (A1) protein along selected transects through representative granules. Overlapping peaks between G3BP1 and A1 indicate their spatial co-enrichment within stress granules. B) Representative confocal images showing colocalization of N protein with G3BP1 in HeLa cells. After fixation, cells were stained with anti-N, and G3BP1, along with DAPI for nuclear staining. N protein shows diffuse cytoplasmic distribution under control conditions; it forms puncta upon stress that overlap with G3BP1-positive stress granules. Merged images show colocalisation of the respective proteins within the cytoplasm. Insets showing magnified views of the boxed regions. Scale bar, 10 µm. Line-scan fluorescence intensity profiles showing normalised signal distributions of G3BP1 and N protein along selected transects through representative granules. Overlapping peaks between G3BP1 and N protein indicate their spatial co-enrichment within stress granules.
